# Pattern Recognition Receptor based Prognostic Biomarkers for predicting Survival of Uterine Corpus Endometrial Cancer Patients

**DOI:** 10.1101/2020.11.25.397703

**Authors:** Dilraj Kaur, Chakit Arora, G.P.S Raghava

## Abstract

In this study, we attempted to identify prognostic biomarkers for predicting survival risk of uterine corpus endometrial cancer (UCEC) patients from the gene expression profile of pattern recognition receptors (PRRs). A wide range of feature selection techniques have been tried, including network-based methods to identify a small number of genes from 331 PRR genes. Firstly, a risk stratification model has been developed using biomarker genes selected using a network-based approach and achieved HR=1.37 with p=0.294. Secondly, we developed a risk stratification model using biomarker of seven genes obtained from clustering and achieved HR=9.14 and p= 1.49×10^-12^. Finally, we developed various combinatorial models using biomarker of 15 PRR genes that were significantly associated with UCEC survival. We found that a multiple genes-based risk stratification model using nine genes (CLEC1B, CLEC3A, IRF7, CTSB, FCN1, RIPK2, NLRP10, NLRP9 and SARM1) gave the best result (HR=10.70, p=1.1×10^-12^, C=0.76, log-rank-p=8.15×10^-14^). The performance of this model improved significantly when we used the clinical stage of patients in combination with the expression of nine genes and achieved HR=15.23 (p=2.21×10^-7^, C=0.78, log-rank-p=2.76×10^-17^). We also developed classification models that can classify high-risk patients (survive ≤ 4.3 years) and low-risk patients (survive > 4.3 years) and achieved AUROC of 0.86. It was observed that specific genes are positively correlated with overall survival of UCEC patients. Based on these observations, we identified potential immunotherapeutic agents for treating UCEC patients.

## Introduction

In today’s world, Uterine corpus endometrial carcinoma (UCEC) or endometrial cancer (EC) is the sixth most common malignancy in females. Out of the total reported 65,620 cases, 12,590 deaths were estimated for 2020 [1]. In contrast to the declining trends for many common cancers, mortality has remained roughly the same for endometrial cancer [2]. Through advancement in high throughput technology, prognosis, and diagnosis of endometrial cancer at an early stage is more often possible, still a notable proportion of patients who develop metastasis or recurrent tumour have unfavorable prognosis. Endometrial cancer is classified under two major subtypes. The first one is the type I tumours, which represent about 75% to 80% of the pathologic subtype and are endometrioid adenocarcinomas [3,4] Whereas type II tumours are serous carcinoma and are less well-differentiated, thus have poorer prognosis [4,5]. Poor diagnosis and prognostic factors lead to high mortality and recurrence in UCEC patients. Till date, a multi-step diagnosis process which includes gynaecological examination, transvaginal ultrasonography (TVUS) and an endometrial biopsy is required to distinguish UCEC from other benign diseases. Clinical features (like tumour grade, cervical involvement, lymph node status, histological subtype) are commonly used as prognostic factors for UCEC patients [6]. The information related to tumour grade, histological subtype and other clinical features is obtained from the biopsies. Though platinum-based chemotherapy and hormonal therapy are the first-line therapies in the case of UCEC, standard methods of treating UCEC consist of primary hysterectomy and bilateral salpingo-oophorectomy [7]. However, it has been observed that due to distant metastasis, patients respond poorly to conventional therapies and have a very low 5-year survival rate (~17%) [8]. Thus, efficient risk stratification methods are required for prognostic evaluation and therapeutic decision making in UCEC patients.

Due to advent of high-throughput sequencing methods, many genetic biomarkers have been identified for diagnosis, classification, and prognosis of UCEC. These biomarkers, in contrast to clinicopathological factors, are related to underlying molecular mechanisms and offer robust theories for the pathogenesis of endometrial cancer. Previous studies revealed that a set of five genes (BUB1B, CCNB1, CDC20, DLGAPS, NCAPG) is helpful in the prognosis of endometrial I carcinoma [9]. Nine genes associated with glycolysis (CLDN9, B4GALT1, GMPPB, B4GALT4, AK4, CHST6, PC, GPC1, and SRD5A3) were shown to be related to overall survival. Amongst these genes, GMPPB had the highest HR of 1.544 and p-value of 0.0134 [10]. Higher KLHL14 expression was associated with worse overall survival (p=0.0370) and progression-free survival (p=0.081) in UCEC samples [11]. Yet another study showed that the TME composition affects the clinical outcomes of UCEC patients, and genes involved are TMEM150B, SIGLEC1, and CTSW. The expression of TMEM150B, CACNA2D2, TRPM5, NOL4, CTSW, and SIGLEC1 were seen to be significantly correlated with overall survival times of UCEC patients [8].

The pattern recognition receptors (PRRs) are known for many years for their role in recognizing microbial ligands and the subsequent activation of the immune system. Recent advancement in the field of bioinformatics led to the development of specific database related to pattern recognition receptor [12]. Recently updated PRRDB2.0 [13] covers the latest information about receptor and their associated ligands suggest that ligands for the toll-like receptors (TLRs), a well-known family of PRRs, shows antitumoral activities in several cancers through activation in tumour cells. This activation could trigger both pro- or antitumoral effects depending on the context [14]. Increased angiogenesis and survival, enhancement of tumour invasion, resistance to apoptosis are some of the functions which show that TLRs act as tumour promoters [15,16]. The TLR pathways are critical regulators in chemo-resistance, possibly through activated NF-kB, upregulated expression of the antiapoptotic protein Bcl-2 [17]. Toll-like receptors like TLR3, 4, 7 and 9 expression correlates with poor differentiation, high proliferation, advanced staging in oesophageal cancer [18,19]. In lung cancer, TLR5 expression is connected with a good prognosis, whereas TLR7 with poor diagnosis. Also, TLR4, 5, 8 and 9 expressions are found to be higher in lung cancer patients than in healthy cancer tissue [20]. TLR7 and 8 are overexpressed and stage-dependent in case of pancreatic cancer as compared to healthy pancreas [21]. TLR1 expression in the favourable prognosis of pancreatic cancer (HR = 0.68, 95% CI 0.47-0.99, p = 0.044) [22]. TLR9 expression is associated with good survival in Renal cell carcinoma [23] is elevated in glioblastoma stem cells [24] and its lower expression is helpful in prediction of disease-free survival in the case of triple-negative breast cancer [25]. In the case of epithelial ovarian cancer, a study showed the association of high expression of TLR4 and MyD88 with poorer overall survival in patients (HR 2.1, 95 % CI 1.1-3.8) [26]. Previously reported studies show the variant alleles of the TLR9 polymorphisms, rs5743836 (C allele) and rs187084 (C allele) combined, can be associated with a decreased risk in the case of UCEC (OR 0.11, 95% CI 0.03-0.44, p = 0.002). The high activity C allele of rs5743836 may allow a better response to pathogenic microbes that are present in the endometrium [27]. It also provides chemo-resistance in ovarian cancer cells [28]. TLR3 and TLR4 expression were examined during the menstrual cycle, endometriosis, postmenopausal endometrium, endometrial hyperplasia and low expression of TLR3 and 4 were shown to be associated with poor prognosis in UCEC [29].

In this study, we exploited the mRNA expression data obtained from The Cancer Genome Atlas-Uterine Corpus Endometrial Carcinoma (TCGA-UCEC) cohort. Firstly we estimated the prognostic performance of each PRR gene by means of survival analysis. We used gene coexpression network-based feature selection and clustering-based approach to find critical PRR genes. Next, we created risk stratification models using these obtained genes. Further, to improve the performance of the model, we identified 15 critical PRRs related genes that are associated with UCEC prognosis viz. CLEC1B, CLEC3A, MRC1, IRF7, CTSB, FCN1, RIPK2, CLEC3B, CLEC12B, TLR4, NLRP10, NLRP9, TNIP1, SARM1 and MAPKAPK2 based on cox-regression survival analysis. The 9 gene (CLEC1B, CLEC3A, IRF7, CTSB, FCN1, RIPK2, NLRP10, NLRP9 and SARM1) voting based model was found to perform the best and also stratified high-risk clinical groups significantly. Finally, after a comprehensive prognostic comparison with other clinicopathological factors, we developed a hybrid model which combines expression profile of nine genes with ‘Clinical stage’ to predict High and Low-risk UCEC patients with high precision. We also excerpted candidate biomolecules that can help modulate the gene expressions and also possibly act as drugs in the treatment of UCEC patients.

## 2 Materials and Methods

### 2.1 Dataset and pre-processing

The dataset was originally taken from ‘The Cancer Genome Atlas’ (TCGA) using TCGA Assembler 2 [30] which consisted of quantile normalized RNA-seq expression values for 201 Uterine Corpus Endometrial Carcinoma (UCEC) patients. The dataset had information about overall survival (OS) and censoring for only 541 patients. The list of 331 pattern recognition receptor signaling pathway genes was taken from GSEA [31] and HGNC [32]. Out of which gene expression data was available only for 308 PRR genes. Thus, the final dataset was reduced to 541 samples constituting RNA-seq values for 308 PRRs related genes using in-house python and R scripts. We have summarized clinical, demographic and pathologic features corresponding to the UCEC final dataset in Supplementary S1 Table 1.

### 2.2 Survival analysis

Hazard ratios (HR) along with confidence intervals (95% CI) were computed to predict the risks of death associated with high-risk and low-risk groups based on overall survival time of patients. These were stratified based on appropriate cut-offs for various factors, using the univariate unadjusted Cox-Proportional Hazard (Cox-PH) regression models. Kaplan-Meier (KM) plots were used to compare survival curves of high risk and low-risk groups. Survival analyses on these datasets were performed using ‘survival’ and ‘survminer’ packages (V.2.42-6) in R (V.3.4.4, The R Foundation). Statistical significance between the survival curves was estimated using log-rank tests. Wald tests were performed to evaluate the importance of the explanatory variables used for HR calculations. Concordance index (C) provided the strength of the predictive ability of the model [33–35] p-values less than 0.05 were considered as significant. Multivariate survival analysis based on Cox regression was employed to compare the relationship between various covariates.

### 2.3 Gene co-expression network

PRR genes co-expression network was constructed to identify the interaction among the genes. To formulate the genes co-expression network used here, we applied Pearson correlation (R) for each PRR genes pair using gene expression value, to calculate statistically significant associations. Igraph’ package in R was utilized to create an edge list from the obtained matrix using correlation cutoff, |R| >0.5. After that, a gene co-expression network was constructed using Cytoscape 3.7.1. ‘Network analyzer’ from Cytoscape was used to evaluate the degree of each node.

### 2.4 k-means and k-medoids clustering

k-means unsupervised clustering was used to identify a pre-specified number (k) of representatives/centroids. Package ‘sklearn’ was used to partition all PRR genes into k clusters in which each PRR gene belongs to the respective cluster with the nearest mean. k-means clustering was done using Euclidean distance between gene expressions of 308 PRR genes. After k clusters are formed, a representative gene from each cluster is chosen. This choice is made based on the least p-value based on univariate survival analysis. Similarly, medoids from each cluster were selected as representatives. This is due to the fact that unlike centroids, medoids are always restricted to be genes of the clusters. Further, we used obtained representative genes for model development.

### 2.5 Multiple gene based models

#### 2.5.1 Machine learning based regression (MLR) models

Regression models from ‘sklearn’ package in Python were implemented to fit the gene expression values (independent variables) against the OS time (target variable). Various regressors such as Linear, Random-forest (RF), K-nearest neighbours (KNN), Ridge, Lasso, Lasso-Lars and Elastic-Net were used. The fitting and test evaluations were carried using a five-fold cross-validation scheme, as implemented in previous studies [36–38]. Combination of all five evaluated test datasets (predicted OS) was then used to classify the actual patient survival time (OS) at median cut-off to estimate HR, CI and p-values. Hyperparameter optimization and regularization was achieved using the in-built function ‘GridsearchCV. Model’s performance is denoted using standard parameters viz. RMSE (root mean squared error) and MAE (mean absolute error).

#### 2.5.2 Prognostic Index (PI)

As implemented in [39,40] PI for a set of *k* genes was evaluated as:

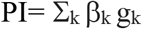

Where, *β* represents regression coefficient obtained for a gene *g* as estimated from a univariate Cox regression. PI for different set of genes was used for stratifying risk groups and standard metrics such as HR, p-value etc. were estimated.

#### 2.5.3 Gene voting based model

Corresponding to an individual gene expression (median cut-off), a risk label ‘High Risk’ or ‘Low Risk’ was assigned to each patient. Thus for *n* survival associated genes, every patient was denoted by a vector of *n* risk labels. In gene voting based method, the patient is ultimately classified into one of the high/low risk categories based on the dominant ‘label’ (i.e. occurring more than n/2 times) in this vector. This is followed by evaluation of standard metrics.

## 3. Results

### 3.1 Survival associated Pattern recognition receptor (PRRs) genes

A univariate Cox-PH analysis was done for all 308 PRR genes using median expression cutoffs. The results of survival analysis in the form of parameters HR and p-values are provided in Supplementary S1 Table 2. Out of 308 genes, only fifteen genes were found to be significant (p <0.05). The survival analysis revealed a total of 9 good prognostic marker (GPM) genes i.e the genes that are positively correlated with patient OS time and 6 bad prognostic marker (BPM) genes which are negatively correlated with OS time of the patients. Table 1 shows the results for these genes along with the metrics associated with stratification of high/low risk patients. The KM plots are shown in Supplementary S2 Figure 1. The precise molecular information about these 15 genes and PMIDs of the studies pertaining to their role in cancer, as obtained from Gene cards [41] and The Candidate Cancer Gene Database (CCGD) [42] respectively, is provided in Supplementary S1 Table 3. Supplementary S1 Table 4 shows results of risk stratification performed using various previously suggested prognostic genes in UCEC using cox univariate analysis in TCGA-UCEC dataset at median expression cutoff.

**Table 1.** The table shows results of univariate cox regression with > median cut-off. Genes with HR>1 are BPM while HR<1 are GPM.

| Gene | HR | 1/HR | p-value | C-INDEX | 95%CI(L) | 95%CI(U) | log rank (p) |
| --- | --- | --- | --- | --- | --- | --- | --- |
| CLEC1B | 6.48 | 0.15 | 2.11E-06 | 0.58 | 2.99 | 14.04 | 1.01E-04 |
| CLEC3A | 2.71 | 0.37 | 3.47E-03 | 0.58 | 1.39 | 5.28 | 7.16E-03 |
| MRC1 | 2.17 | 0.46 | 1.60E-02 | 0.60 | 1.16 | 4.09 | 1.23E-02 |
| IRF7 | 0.47 | 2.14 | 1.77E-02 | 0.59 | 0.25 | 0.88 | 1.43E-02 |
| CTSB | 0.50 | 2.00 | 2.65E-02 | 0.61 | 0.27 | 0.92 | 2.31E-02 |
| FCN1 | 2.00 | 0.50 | 2.85E-02 | 0.56 | 1.08 | 3.72 | 2.47E-02 |
| RIPK2 | 0.50 | 2.00 | 2.86E-02 | 0.57 | 0.27 | 0.93 | 2.42E-02 |
| CLEC3B | 0.51 | 1.94 | 3.35E-02 | 0.59 | 0.28 | 0.95 | 2.96E-02 |
| CLEC12B | 1.86 | 0.54 | 3.82E-02 | 0.57 | 1.03 | 3.35 | 4.11E-02 |
| TLR4 | 0.53 | 1.89 | 3.88E-02 | 0.55 | 0.29 | 0.97 | 3.55E-02 |
| NLRP10 | 1.94 | 0.52 | 3.99E-02 | 0.57 | 1.03 | 3.65 | 5.00E-02 |
| NLRP9 | 0.53 | 1.89 | 4.15E-02 | 0.56 | 0.29 | 0.98 | 3.70E-02 |
| MAPKAPK2 | 0.53 | 1.88 | 4.35E-02 | 0.55 | 0.29 | 0.98 | 3.89E-02 |
| TNIP1 | 0.54 | 1.86 | 4.38E-02 | 0.56 | 0.29 | 0.98 | 4.02E-02 |
| SARM1 | 0.54 | 1.85 | 4.95E-02 | 0.55 | 0.29 | 1.00 | 4.54E-02 |

### 3.2 Risk prediction using network-based features

In this section, an attempt has been made to select features from network of PRR genes. The aim of using network to identify features/representatives was to understand connectivity amongst PRR genes. We used following approaches for feature selection: i) Hub genes of the network, ii) Medoids of clusters and iii) Representatives of clusters. These selected features or PRR genes have been used to develop models for predicting survival of cancer patients.

#### 3.2.1 Hub genes of network

A correlation matrix was computed from PRR genes where correlation among all possible pair of genes was computed based on expression. This correlation matrix was used to create edges of network using Igraph for highly correlated pair of genes as described in Material & Methods. We used ‘Cytoscape’ software to visualise and analyse the gene network. The effective correlation was set to be greater than 0.5, which resulted in 116 nodes and 804 edges. We selected top 15 hub genes (viz. BTK, ITGB2, HAVCR2, FCRL3, CD163, CD300LF, CD68, CTSS, CLEC10A, CLEC12A, NR1H3, CLEC4E, CD209, ITGAM and TLR8) based on their degree. These hub genes were used to build prognostic model for predicting survival risk UCEC patients. Our voting based model developed using these 15 hub genes achieved HR=1.37 with p=0.294. The network is shown in Figure 1. Also, the network metrics are provided in Supplementary S1 Table 5.

**Figure 1.**
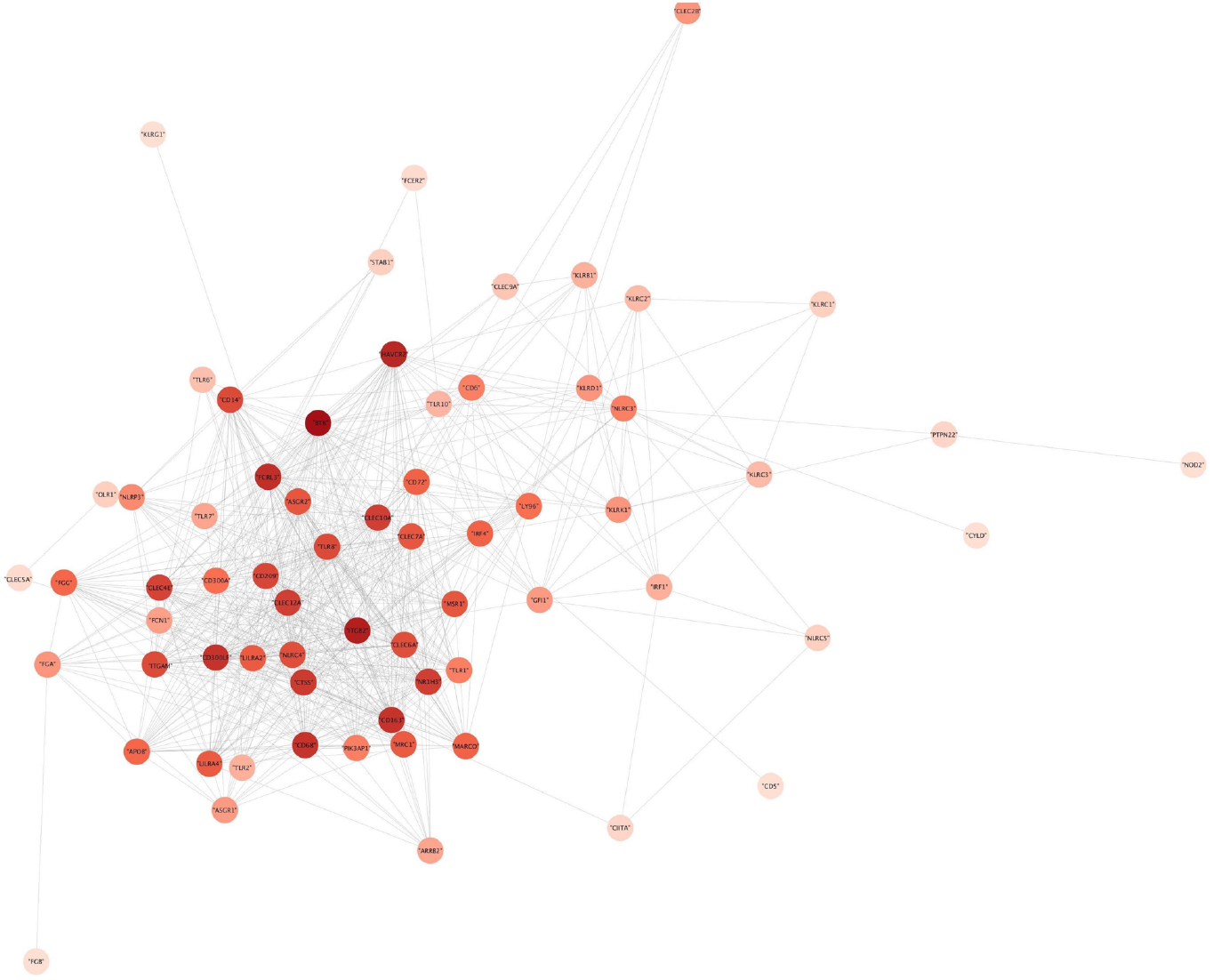
Gene co-expression network analysis of PRR genes. Interconnection of 15 hub genes; darker colour represents a higher degree score

#### 3.2.2 Medoids of clusters

We performed k-medoids clustering of genes based on pairwise dissimilarity. The medoids were selected for clusters at different k = 5,10,15,20 and 25. Best result was obtained for gene voting model at k=5 (HR=1.85, P=0.045). The results at different k values is provided in Supplementary S1. Table 6.

#### 3.2.3 Representatives of clusters

The representative gene, from each cluster, was chosen on the basis of least p-value obtained from univariate survival analysis. Obtained representative genes from each cluster were used to develop risk stratification model. This process was repeated for k = 5, 10, 15, 20 and 25. Best result was obtained for k=10 (HR= 4.11, p= 3.7×10^-5^) as shown in Table 2.

**Table 2.**
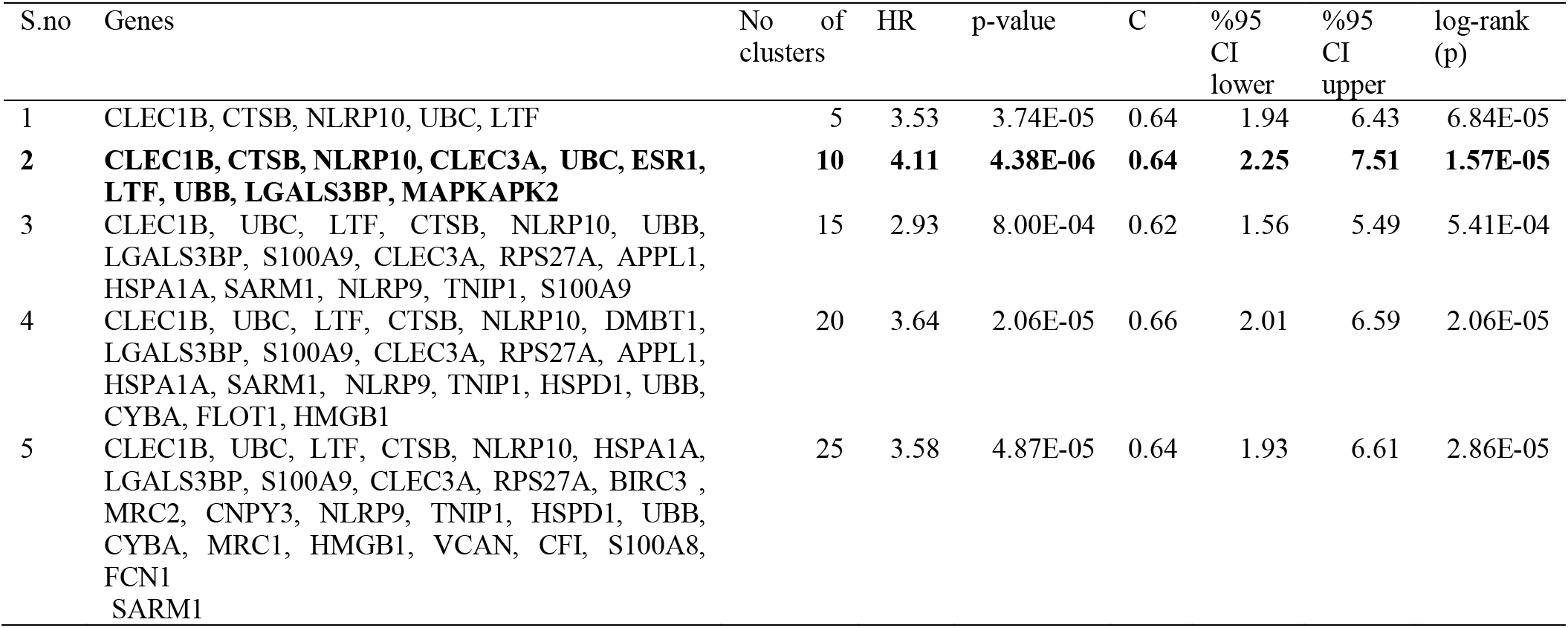
The table shows results of gene voting model for chosen representative genes from each cluster.

Further, we filtered the representative genes by using a cut-off of p <0.05 within each cluster. Best results were obtained for gene voting models as shown in Table 3. It was observed that optimal risk segregation was obtained for k=15 which resulted in 7 representative genes. The detail about distribution of 308 genes into 15 clusters is provided in Supplementary S1 Table 7. The gene voting model developed for these genes (CLEC1B, CLEC3A, CTSB, NLRP10, NLRP9, TNIP1 and SARM1) achieved HR=9.14 and p-val= 1.49×10^-12^. The network construction is done by ‘cytoscape 3.7.1’ and shows 7 different clusters with their representative genes in Figure 2.

**Table 3.** The table shows results of gene voting model for chosen significant representative genes only from each cluster.

| S.no | Representative genes | No of clusters | HR | p-value | C | %95 CI lower | %95 CI upper | log-rank (p) |
| --- | --- | --- | --- | --- | --- | --- | --- | --- |
| 1 | CLEC1B, CTSB, NLRP10 | 5 | 3.62 | 1.12E-04 | 0.60 | 1.88 | 6.96 | 4.87E-04 |
| 2 | CLEC1B, CTSB, NLRP10, CLEC3A | 10 | 4.25 | 1.63E-06 | 0.53 | 1.30 | 13.84 | 4.70E-02 |
| 3 | **CLEC1B, CLEC3A, CTSB, NLRP10, NLRP9, TNIP1, SARM1** | **15** | **9.14** | **1.49E-12** | **0.73** | **4.95** | **16.87** | **6.64E-12** |
| 4 | CLEC1B, CLEC3A, CTSB, NLRP10, NLRP9, TNIP1, SARM1 | 20 | 9.14 | 1.49E-12 | 0.73 | 4.95 | 16.87 | 6.64E-12 |
| 5 | CLEC1B, CLEC3A, CTSB, NLRP10, NLRP9, TNIP1, SARM1, MRC1, FCN1 | 25 | 5.46 | 3.05E-08 | 0.67 | 2.99 | 9.95 | 9.00E-08 |

**Figure 2.**
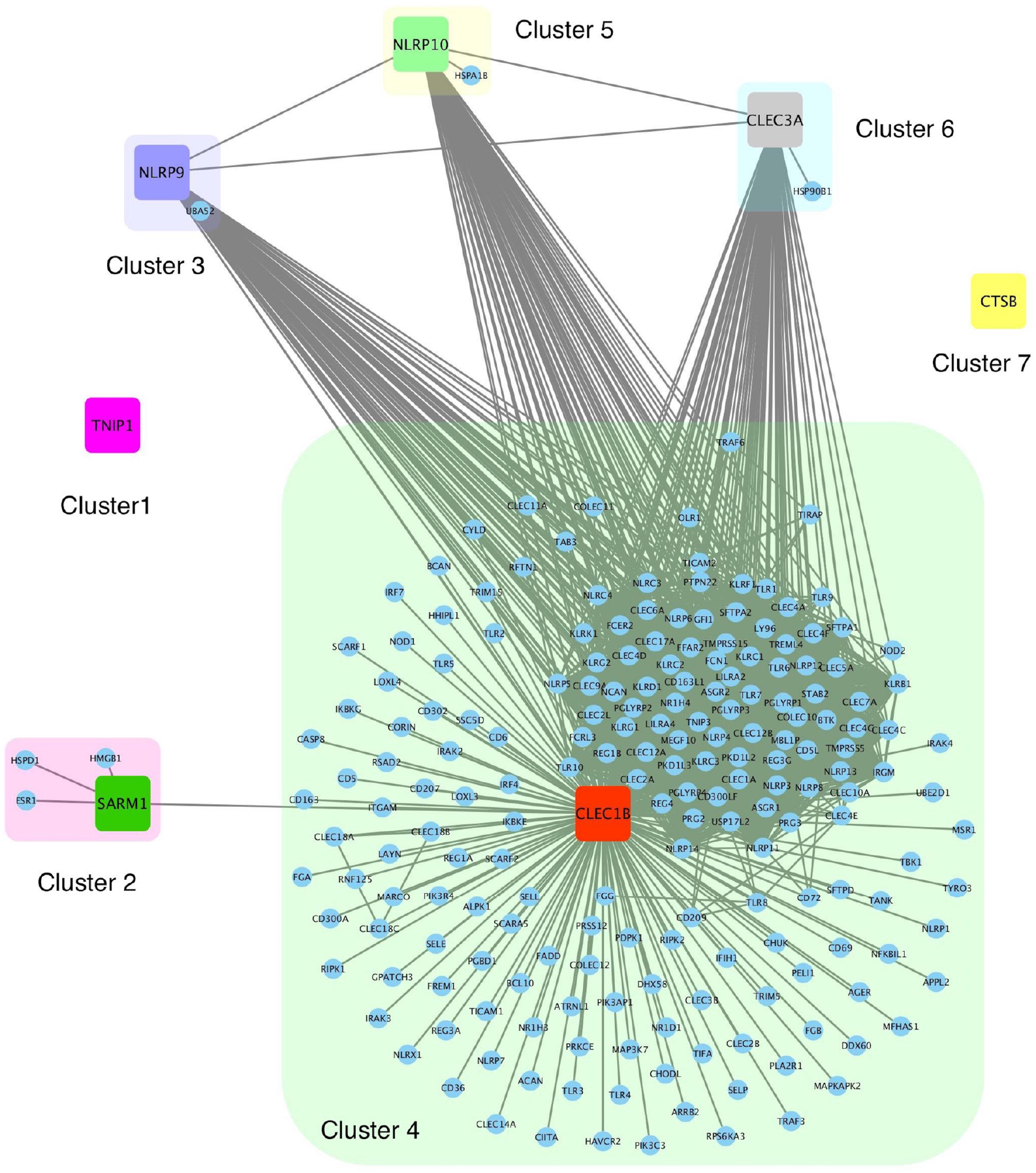
Network based on clustering: 7 different clusters are shown in different colors. Representative PRR genes are highlighted and represented within large squares.

### 3.3 Risk estimation using multiple gene-based models

Several risk stratification models based on MLR, prognostic index and gene voting were constructed using the expression profile of survival associated PRR genes (based on p-value). We tried various combination using 15 significant genes for this. It was found that combination of nine genes i.e. CLEC1B, CLEC3A, IRF7, CTSB, FCN1, RIPK2, NLRP10, NLRP9 and SARM1 performed the best. Table 4 shows the results corresponding to various risk prediction models for these nine genes. Amongst these, the performance of gene voting based model was found to be the best with HR=10.70 and p~10^-12^. A C value of 0.76 was also the highest for this model and high/low risk groups survival curves were significantly separated with a logrank-p~10^-14^. Figure 3 shows the KM plot representing the survival curves corresponding to the two risk groups. While the 5-year survival rate for low risk patients was close to 85%, for high risk patients it was seen to drop as low as 15%. PI based model performed the second best with HR=3.41 and p~10^-3^ and regression based Linear model was the third best (and top amongst MLR models) with HR=1.66 but p-value was found to be statistically insignificant.

**Table 4.**
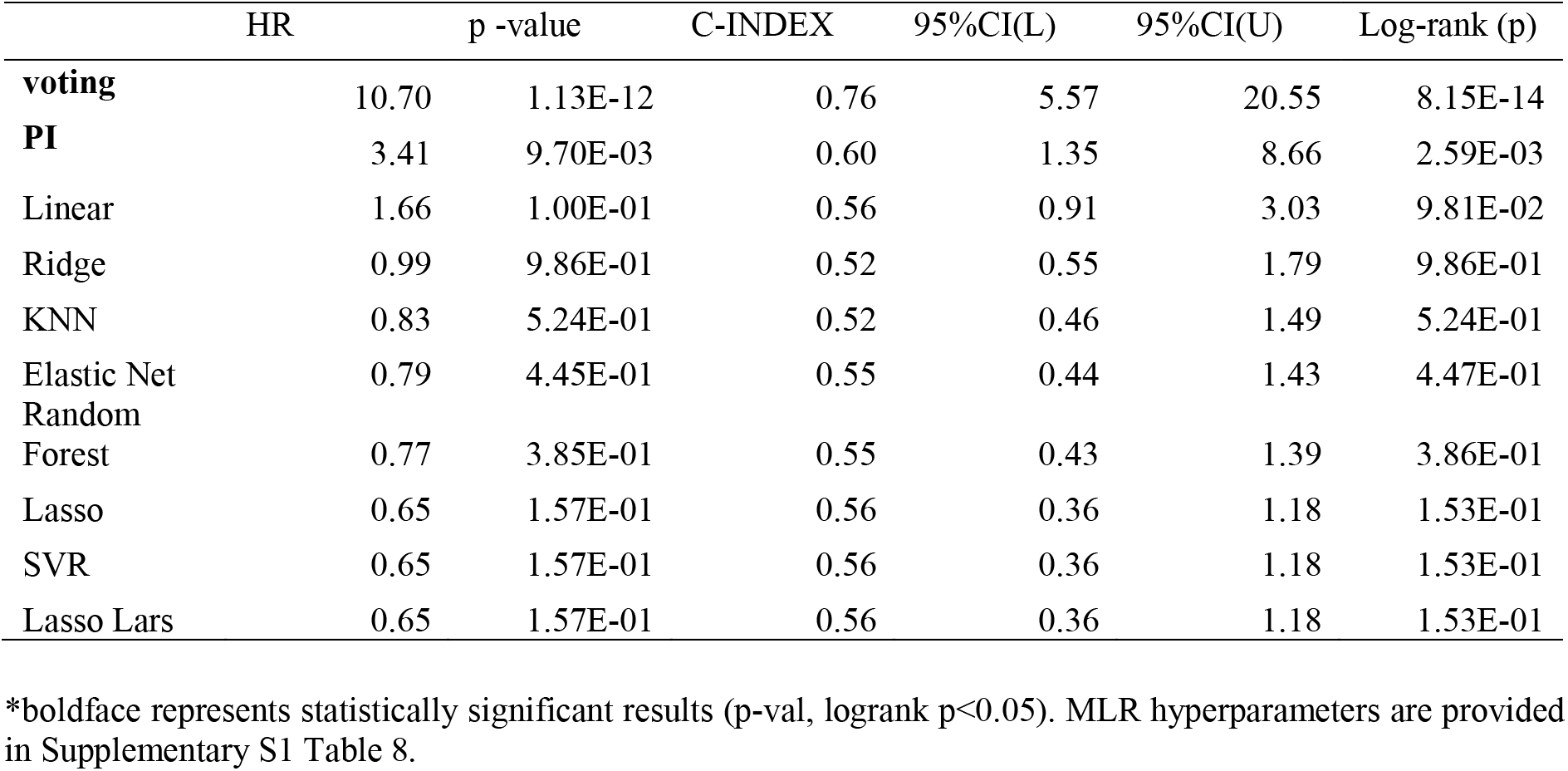
The performance of different models developed using multiple gene expression profile.

|  | HR | p -value | C-INDEX | 95%CI(L) | 95%CI(U) | Log-rank (p) |
| --- | --- | --- | --- | --- | --- | --- |
| **voting** | 10.70 | 1.13E-12 | 0.76 | 5.57 | 20.55 | 8.15E-14 |
| **PI** | 3.41 | 9.70E-03 | 0.60 | 1.35 | 8.66 | 2.59E-03 |
| Linear | 1.66 | 1.00E-01 | 0.56 | 0.91 | 3.03 | 9.81E-02 |
| Ridge | 0.99 | 9.86E-01 | 0.52 | 0.55 | 1.79 | 9.86E-01 |
| KNN | 0.83 | 5.24E-01 | 0.52 | 0.46 | 1.49 | 5.24E-01 |
| Elastic Net | 0.79 | 4.45E-01 | 0.55 | 0.44 | 1.43 | 4.47E-01 |
| Random Forest | 0.77 | 3.85E-01 | 0.55 | 0.43 | 1.39 | 3.86E-01 |
| Lasso | 0.65 | 1.57E-01 | 0.56 | 0.36 | 1.18 | 1.53E-01 |
| SVR | 0.65 | 1.57E-01 | 0.56 | 0.36 | 1.18 | 1.53E-01 |
| Lasso Lars | 0.65 | 1.57E-01 | 0.56 | 0.36 | 1.18 | 1.53E-01 |

\*boldface represents statistically significant results (p-val, logrank $p < 0.05$ ). MLR hyperparameters are provided in Supplementary S1 Table 8.

**Figure 3.**
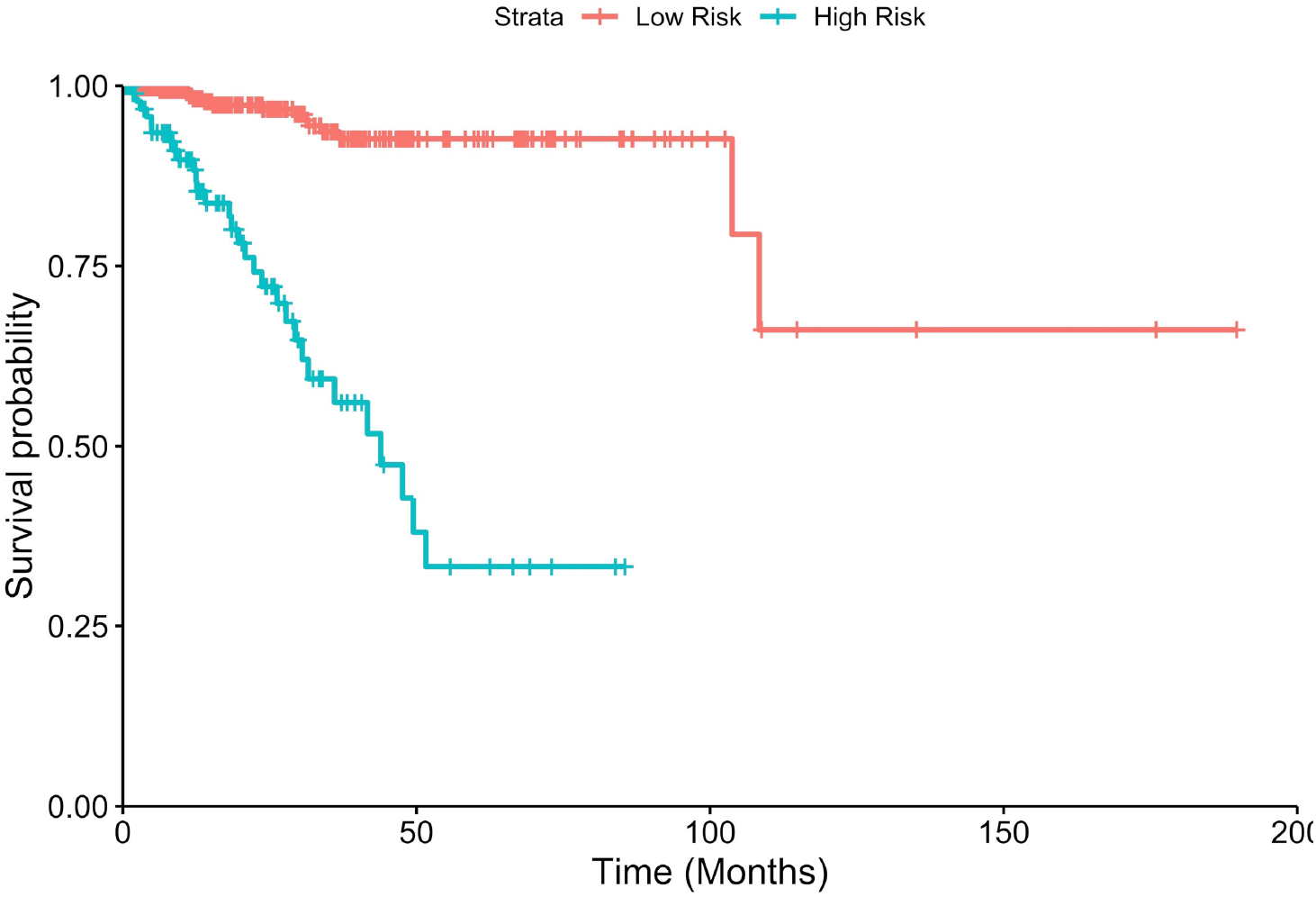
KM plot showing risk stratification of UCEC patients based on gene voting model. Patients with greater than four ‘high risk’ labels in the 10-bit risk vector are assigned (blue) as High Risk (HR=10.70, p=1.13×10^-12^, C=0.76, log-rank-p =8.15×10^-14^) while others were assigned as Low Risk (red).

### 3.4 Multiple gene model sub-stratifies patients in clinico-pathological high-risk groups

In the past, several studies indicate the role of certain clinico-pathological factors in UCEC prognosis such as Histologic diagnosis, ethnicity, clinical stage [6] menopause status, peritoneal washing etc [43]. Thus, we performed a univariate analysis to assess the association of these factors with OS in our dataset. Table 5 shows the results of the univariate analysis. The clinical factors like clinical stage, residual tumor, peritoneal washing, grade, histologic grade, menopause status are seen to be the significant factors in the UCEC prognosis.

**Table 5.** Univariate analysis using clinico-pathological features. Clinical stage, residual tumor, peritoneal washing, grade, histologic washing, menopause status are seen to be the significant factor.

| Clinical Factors | Strata | n | HR | p-value | C-INDEX | 95%CI(L) | 95%CI(U) | log-rank (p) |
| --- | --- | --- | --- | --- | --- | --- | --- | --- |
| **clinical stage** | III, IV vs I,II | 541.00 | 4.44 | 1.31E-06 | 0.71 | 2.43 | 8.11 | 8.81E-07 |
| **Residual tumor** | R0 vs R1,R2 | 411.00 | 0.29 | 8.06E-04 | 0.59 | 0.14 | 0.60 | 2.27E-03 |
| **Peritoneal washing** | negative vs positive | 407.00 | 0.31 | 2.42E-03 | 0.58 | 0.15 | 0.66 | 5.38E-03 |
| **Grade** | G3, High grade vs G1,G2 | 541.00 | 3.27 | 2.46E-03 | 0.61 | 1.52 | 7.03 | 7.24E-04 |
| **Histologic_diagnosis** | MSE, SEA VS EEC | 541.00 | 2.29 | 6.61E-03 | 0.56 | 1.26 | 4.16 | 8.65E-03 |
| **Menopause status** | Pre vs post | 512.00 | 2.54 | 3.51E-02 | 0.53 | 1.07 | 6.02 | 5.92E-02 |
| age | >64 vs <=64 | 541.00 | 1.30 | 3.79E-01 | 0.54 | 0.72 | 2.33 | 3.79E-01 |
| race | White vs others | 511.00 | 1.15 | 6.83E-01 | 0.50 | 0.58 | 2.29 | 6.79E-01 |
| Ethnicity | Hispanic or latino vs Not hispanic not latino | 388.00 | 1.44 | 7.23E-01 | 0.50 | 0.19 | 10.61 | 7.37E-01 |
| History other malignancy | No vs Yes | 541.00 | 1.18 | 7.84E-01 | 0.49 | 0.36 | 3.81 | 7.79E-01 |
| Surgical approach | open vs minimally invasive | 519.00 | 1.06 | 8.71E-01 | 0.51 | 0.54 | 2.07 | 8.71E-01 |

\*boldface represents statistically significant results (p-val, logrank p<0.05)
\*EEA- Endometrioid endometrial adenocarcinoma, SEA- Serous endometrial adenocarcinoma, MSE- Mixed serous and endometrioid

The gene voting model was able to sub stratify the high risk UCEC patients based on clinicopathological factors like histologic diagnosis, peritoneal washing, menopause status, neoplasmic grade, residual tumor and clinical stage as shown in Figure 4. The KM plots alongwith low logrank-p values denote a significant separation between high and low risk patients.

**Figure 4.**
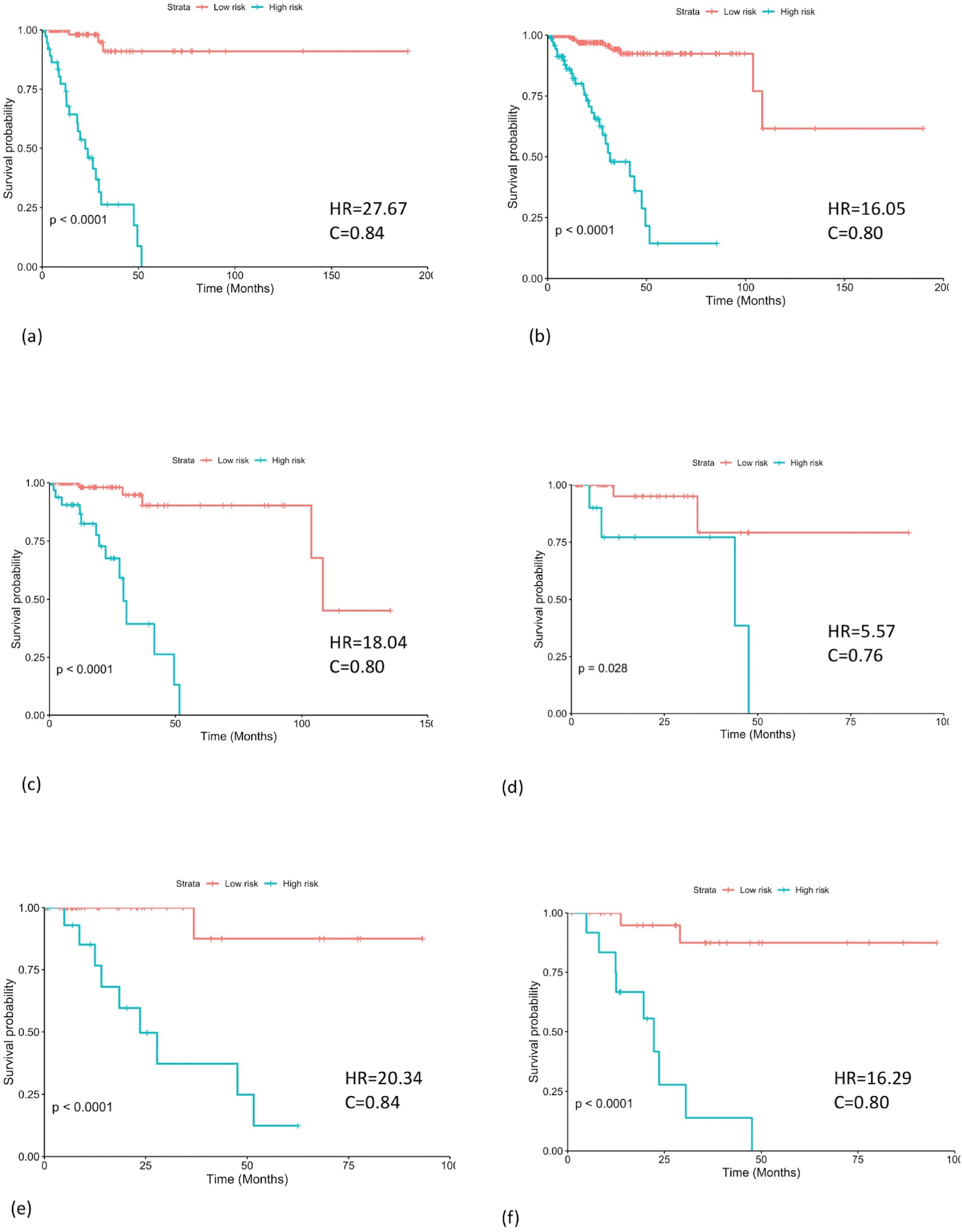
Gene voting model sub-stratifies high risk groups. (a) Patients with clinical stage III/IV (n=152) were stratified into high and low risk groups with logrank-p=4.3×10^-13^ (b) Neoplasmic grade = G3 and High grade (n=321) were stratified into high and low risk groups with logrank-p=5.0×10^-14^ (c) Histologic diagnosis= Mixed serous adenocarcinoma, Endometrioid endometrial adenocarcinoma (n=136) were stratified into high and low risk groups with logrank-p=9.2×10^-8^ (d) Menopause status = pre (n=52) were stratified into high and low risk groups with logrank-p=4.1×10^-9^ (e) Peritoneal washing =positive (n=57) were stratified into high and low risk groups with logrank-p=8.4×10^-5^ (f) Residual tumor = R1,R2 (n=38) were stratified into high and low risk groups with logrank-p=3.2×10^-5^.

### 3.5 Multivariate Analysis

We performed a multivariate cox regression survival analysis using six prominent prognostic markers obtained, i.e. multiple gene voting model, clinical staging, residual tumour, peritoneal washing, histological subtype, menopause status, grade. It can be seen that p-value corresponding to gene voting model (HR=8.17, p<0.001) and clinical stage (HR=3.11, p=0.03) is significant, whereas it is insignificant for others as depicted by forest plot in Figure 5. Thus, an hybrid model can be made using the gene voting model and clinical stage for further improvement in Risk stratification.

**Figure 5.**
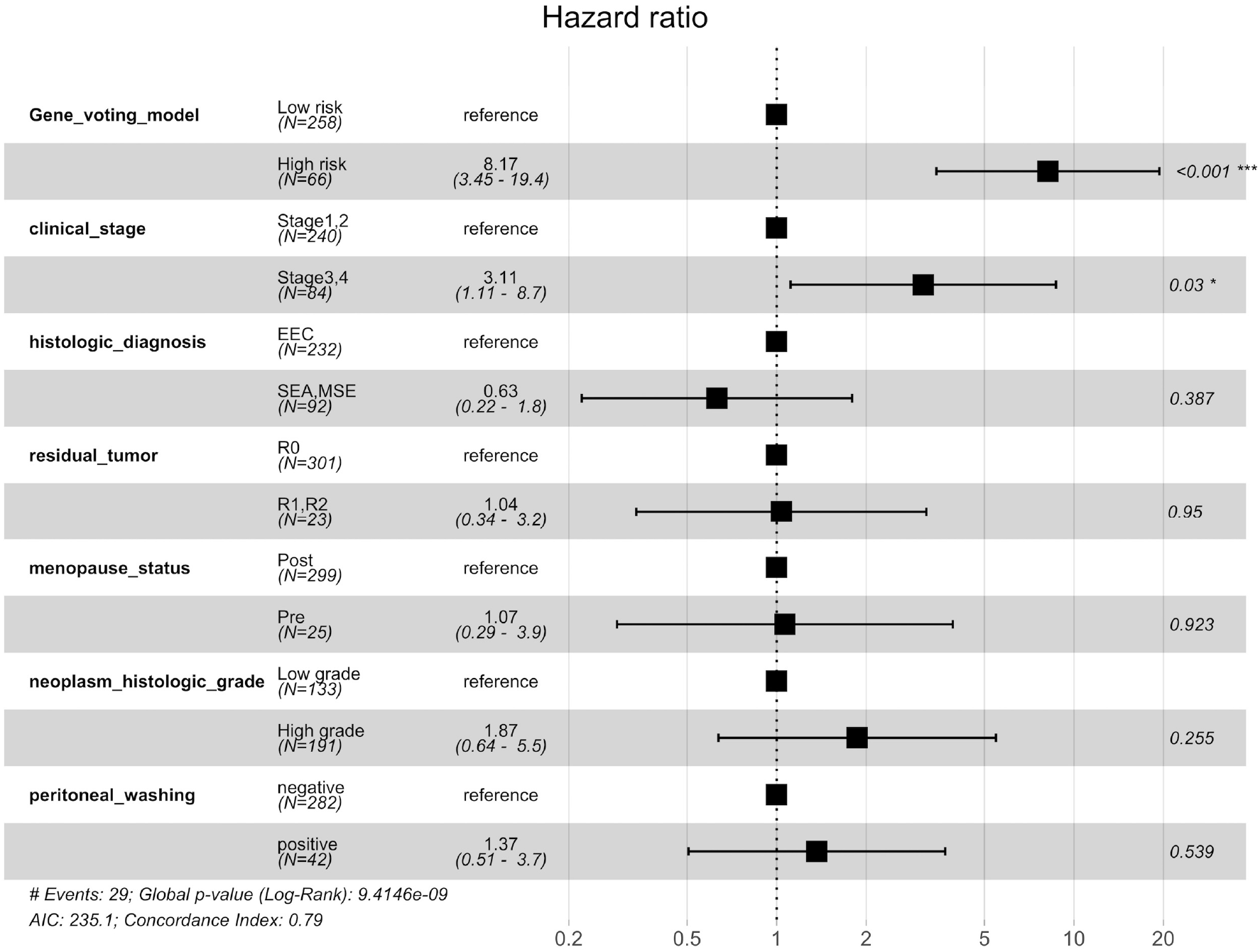
Multivariate analysis reveals gene voting model (HR=8.17, p<0.001) and clinical stage (HR=3.11 p=0.03) as an independent covariates.

### 3.6 Hybrid voting model

After obtaining the independent covariates i.e. multiple gene voting model and clinical stage, based on a multivariate cox regression survival analysis, we developed a hybrid voting model. This model combined clinical stage with the 9-gene voting model for risk stratification purposes. The risk vector associated with each patient was thus now a 10-bit vector with 1 bit assigned to risk label due to clinical stage. We observed that the model performed better than the 9-gene voting model (HR=15.23, p=2.21×10-7, C=0.78, log-rank-p=2.76×10^-17^). Figure 6 shows the KM plot corresponding to the hybrid model.

**Figure 6.**
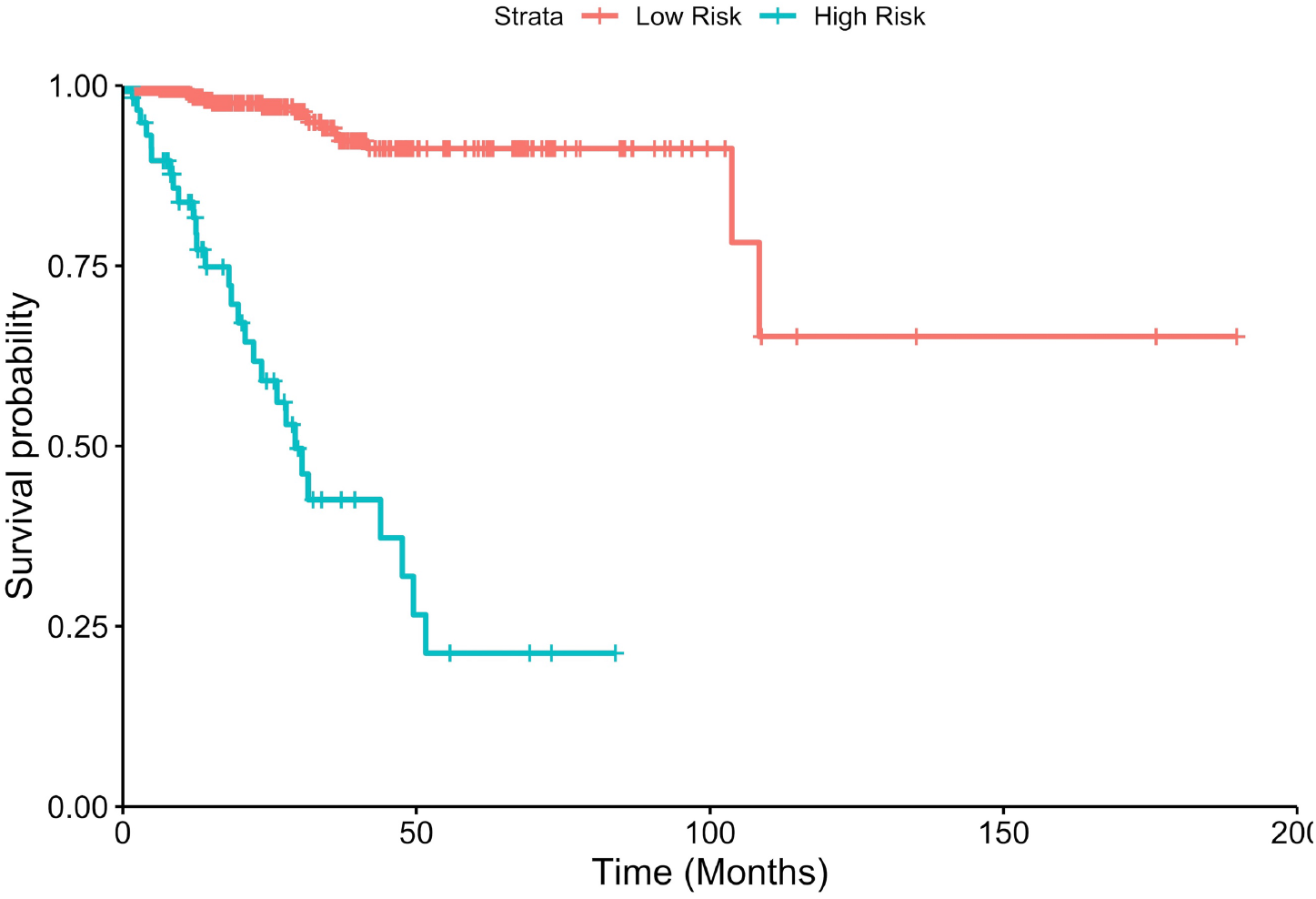
Hybrid model for risk stratification using 9 gene voting model and clinical stage (HR=15.23, p=2.21×10-7, C=0.78, log-rank-p=2.76×10^-17^).

### 3.7 Predictive validation

As implemented in [44] we performed a predictive assessment of our gene voting model using sub-samples of the complete dataset. Sampling sizes of 50%, 70% and 90% were chosen with 100 iterations each. HR and C index were evaluated for each iteration corresponding to the gene voting model and hybrid model. Figure 7 shows the boxplots corresponding to the results. The median HR (15.34, 15.32, 15.02) and C (0.786, 0.792, 0.783) values for hybrid model remain better despite of the sampling size. This method ensured that the risk stratification models were robust and performed well with random datasets of different sizes.

**Figure 7.**
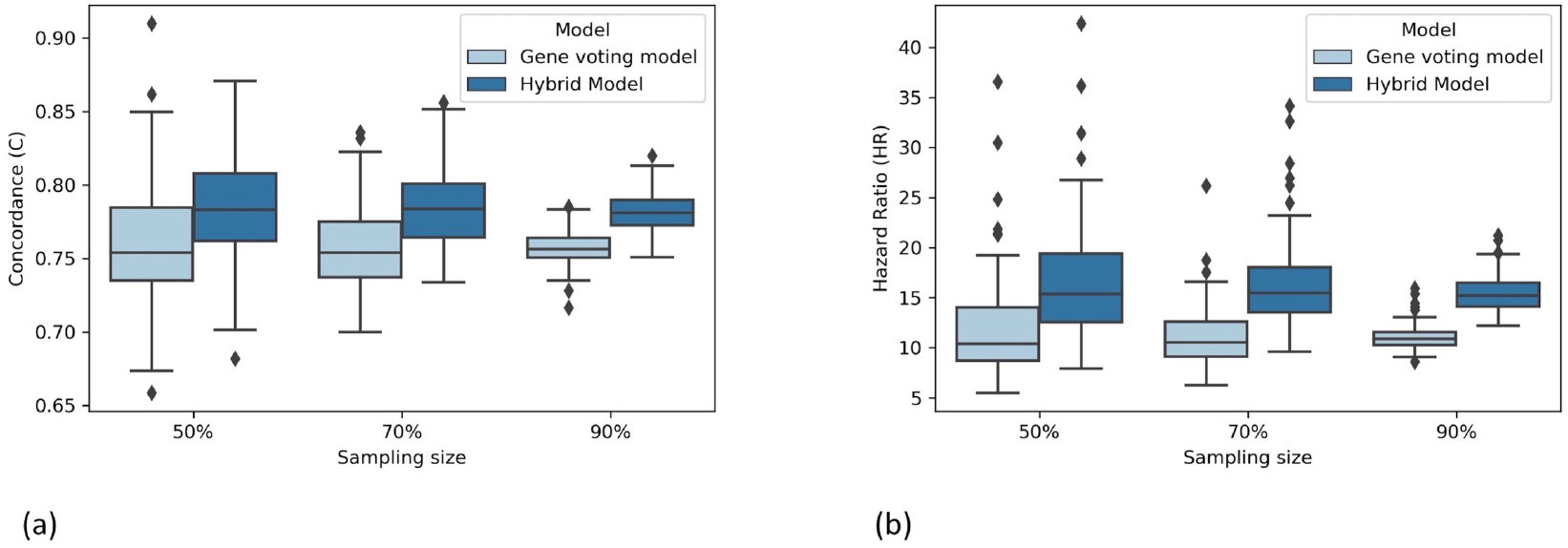
Predictive validation of voting based model (a) Grouped boxplots corresponding to estimated Concordance index (y-axis) for 100 iterations of data sampling (x-axis). (b) Similarly, estimation of Hazard Ratio (y-axis) for different models using random sampling (x-axis).

### 3.8 Classification using hybrid voting model

The performance of hybrid model is evaluated through AUROC (Area under receiver operating characteristic curve) value. ‘survivalROC’ package was used to calculate true positive rate (TPR) and false positive rate (FPR). Here, a true positive prediction being the patient whose OS> cut-off time as well as who was in low-risk group according to the model, while converse applies for a true negative prediction as shown in Figure 8 (a). The median of the OS time was 1.1 yrs. We observed that amongst different OS cutoffs, the model performed best at the cut off of 4.3 years. This cutoff is a good classifier between high and low risk patients. Using this cutoff, an AUROC value of 0.86 was obtained by the classification based on the hybrid voting model. The ROC curve corresponding to this is shown in Figure 8 (b).

**Figure 8.**
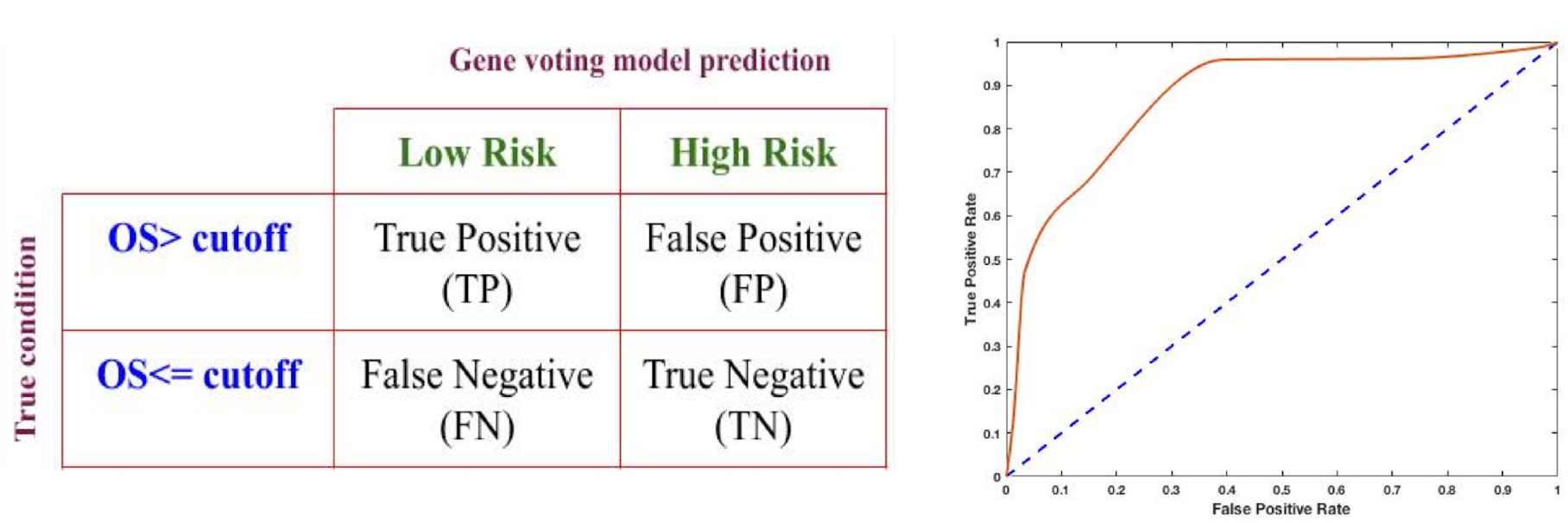
(a) Terminology used for evaluation of confusion matrix. Initial risk labelling was done using an OS cutoff with patients having OS> cutoff labelled as positive or low risk and vice-versa for patients with OS≤cutoff. (b) ROC curve for gene voting model AUROC of 0.86 was obtained.

### 3.9 Screening of therapeutic drug molecules

The choice of therapy is a very crucial step after the identification of genes that play a critical role. Thus, we extracted drug molecules which could re-modulate the overexpressed and underexpressed genes, using the ‘Cmap2 database [45,46] as done in [47]. We queried a list of probe ids corresponding to upregulated genes (CLEC1B, CLEC3A, FCN1, NLRP10) and downregulated genes (RIPK2, SARM1, IRF7, CTSB, NLRP9) as input to Cmap2. The output was ranked based on p-values (results in Supplementary S1 Table 9). Top positive and negative enriched molecules were hexamethonium bromide (enrichment=0.834, p=2.6×10-4) and isoflupredone (enrichment= −0.955, p=2.2 × 10-4) respectively. Hexamethonium bromide is a non-depolarizing ganglion blocker and a (nicotinic acetylcholine receptors) nAChR antagonist. It can be used to treat hypertension and duodenal ulcers [48,49]. It is known to be poorly absorbed from the gastrointestinal tract and does not cross the blood-brain barrier. Isoflupredone, also known as deltafludrocortisone and 9α-fluoroprednisolone, is a synthetic glucocorticoid corticosteroid. Isoflupredone may act as strong inducers to promote the progress of Prostate cancer [50]. 3D conformers of these drug molecules obtained from PubChem [51] are shown in Figure 9.

**Figure 9.**
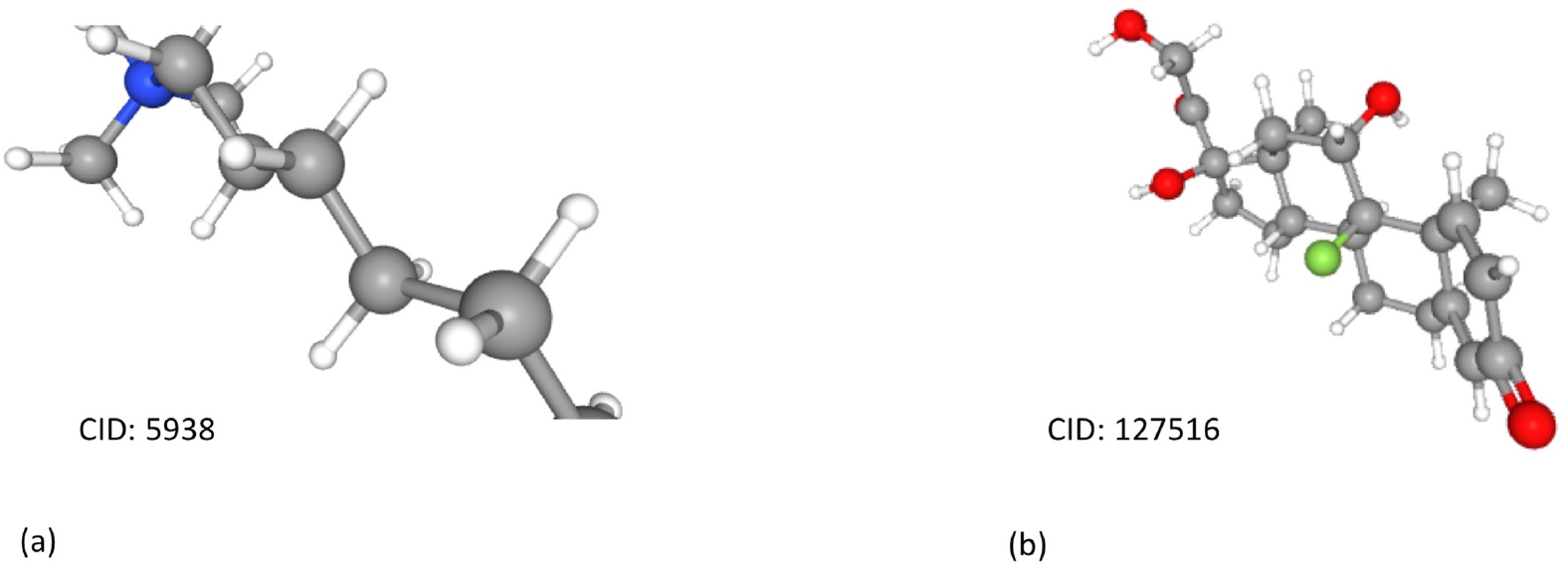
3D conformer of two most significant small molecule drugs (a) Hexamethonium bromide (enrichment=0.834, p=2.6×10^-4^) (b) isoflupredone (enrichment= −0.955, p=2.2 × 10^-4^)

## 4. Discussion

Although UCEC is known to have an excellent prognosis, there remains a decent proportion of patients with an abysmal prognosis. As a result of which, efficient risk assessment strategies are required for clinical decision making and therapeutic intervention. The clinical features like tumour grade, cervical involvement, lymph node status, histological subtype, depth of myometrial invasion, lympho-vascular space invasion (LVSI) that are significant in risk stratification in UCEC patients, are not efficient due to their limitations. Thus, aided by the development in the high-throughput sequencing methods and availability of a massive amount of experimental data, various molecular prognostic markers have been proposed in the past. Earlier studies revealed multiple mechanisms at the molecular level that lead to complicated molecular processes which are crucial for UCEC progression and development. The elucidation of complicated and multifaceted signaling processes orchestrated by PRRs have boosted the therapeutic decision-making strategies in various cancers. Their agonists are used as potentiating systemic treatments such as chemotherapy, targeted therapy and immunotherapy as well as commonly vaccine adjuvants [52]. Amongst numerous examples, agonist/ligands associated to PRRs like MPLA associated with TLR4 were shown to act as potent vaccine adjuvants and promote type 1 T helper (Th1)-biased immune response in case of HPV induced cervical cancer [53,54]. Imiquimod associated with TLR7 was reported to induce apoptosis and stimulate a cell-mediated immune response in the case of basal cell carcinoma [55]. A synthetic derivative of lipid-A, OM-174 associated with TLR2/4 is claimed to reduce tumour progression and prolong survival, especially in combination with cyclophosphamide in the case of melanoma [56]. The ligands/agonists related to TLR4, TL5 and NLR are some of the FDA approved targeted immunotherapy in cervical cancer, various skin cancer and osteosarcoma, respectively. The combination of PRR-based agonist therapy with immune checkpoint-targeted antibodies (such as anti-CTLA-4 or anti-PD-L1 antibodies) may be the future of cancer therapies [52]. However, the role of pattern recognition receptor signalling genes in UCEC is poorly understood.

In the current study, we attempted to examine the expression of PRR genes and their association with UCEC aggressiveness. For this, we evaluated the prognostic potential of these genes in UCEC by employing a recent gene expression dataset. We first used a network-based feature selection for the identification of crucial PRR genes. We calculated the degree of centrality of a gene within the co-expression network, which provided the relative significance of a gene within the network. The higher degree of a gene represented its increased connectivity to other genes and its essential role within the system. However, the performance of the risk stratification model based on 15 hub genes was found to be inferior. Next, to develop a biomarker with minimal genes/features and better prognosis, a clustering approach was implemented. Wherein genes were segregated into different clusters based on similarity/distance between their expression values. After that, representative genes were selected based on various criteria. We observed that the risk prediction model based on representative genes that were better associated with overall survival (p<0.05) performed better. Further, out of the fifteen genes that were significantly associated with UCEC prognosis, combinations of different genes were used to develop models. Amongst these, the performance of gene voting based model based on nine genes was found to be the best. Amongst these nine genes, CLEC1B known as C-type lectin domain family 1member B found very much useful in atherosclerosis disease [57]. Aggretin AACT is a potential candidate for the treatment of tumour metastasis through CLEC-2 blockade [58]. C-Type Lectin-Like Receptor 2 Suppresses AKT Signaling and Invasive Activities of Gastric Cancer Cells by Blocking Expression of Phosphoinositide 3-Kinase Subunits [59]. CLEC3A promotes tumour progression and poor diagnosis in breast invasive ductal cancer. It is a critical therapeutic for breast cancer [60]. CLEC3A, MMP7, and LCN2 known to be associated with the phosphatidylinositol-3-kinase/Akt pathway, as potential novel markers to predict the postoperative recurrence of PNETs [31129920]. Ficolin1 can be considered as supplementary biomarker in acute myeloid leukaemia in adults [p = 0.004, OR = 2.95, 95% CI (1.41-6.16)] [61]. NLRP10 binds to ASC and can inhibit NF-κB activation and apoptosis, as well as caspase-1-mediated IL-1β maturation [62]. SARM mediates intrinsic apoptosis via B cell lymphoma-2 (Bcl-2) family members. SARM suppresses B cell lymphoma-extra large (Bcl-xL) and downregulates extracellular signal-regulated kinase phosphorylation [63]. SARM1 can act as one of the biomarker in colorectal cancer [64]. The overexpression of SARM1 promotes growth and metastasis in Prostate cancer cells [65] targeting RAGE and IRF7 may facilitate vascular repair in diabetes [66] IRF7 regulates the development of granulocytic myeloid-derived suppressor cells through S100A9 trans-repression in cancer [67]. CTSB rs12898G polymorphism CTSB rs12898 is associated with primary hepatic cancer in a Chinese population [68]. NLRP9 act as inflammasome-related molecules a reliable non-invasive tool for Breast cancer diagnosis [69]. The NLRP4 and NLRP9 expressions assessment could be beneficial in predicting the BCG failure and also in decision making for early radical surgery [70].

We also evaluated the risk stratification performance of other genes suggested in the literature and showed that the nine genes proposed in our study show better results (Supplemetary1. Table 4). We also found drug-molecules which could be potentially be used for UCEC therapy and require future efforts. In our study, ‘Hexamethonium bromide’ and ‘isoflupredone’ were two such top molecules found that have anticancer properties and thus used in anticancer treatment. We developed a hybrid model which is a combination of multiple gene expression based voting model using these nine genes and clinical stage. Further, this model was able to discriminate high and low-risk patients among established high-risk groups. We also estimated the performance of the model against significant clinicopathological factors available using multivariate analysis. We performed Monte Carlo validation to check the robustness of the model. The model was also able to achieve an AUROC of 0.86 for the classification of patients having more than 4.3-year overall survival with those having less than or equal to 4.3 years overall survival time. In conclusion, we identified vital PRR genes with a possible role in UCEC pathogenesis and prognosis. While this is supported by previous literature and explored in the current study as an in-silico analysis, it is subjected to further validation. Also, apart from their strong prognostic potential, as elucidated in this study, these genes could also be investigated further in the context of therapeutic targets in UCEC and clinical decision making.

## Author Contributions

DK collected, compiled, and processed the data sets. DK and CA developed computer programs. DK implemented the algorithms and prediction models. DK analyzed the results. DK, CA and GPSR wrote the manuscript. GPSR conceived and coordinated the project and provided overall supervision of the project. All authors have read and approved the final manuscript.

## Acknowledgement

Authors are thankful to J.C. Bose National Fellowship, Department of Science and Technology (DST), and Indraprastha Institute of Information Technology Delhi (IIITD) for fellowships and the financial support.

