## Supplementary S1 for "Pattern Recognition Receptor based Prognostic Biomarkers for predicting Survival of Uterine Corpus Endometrial Cancer Patients"

Figure 1 . KM plot showing risk stratification of UCEC patients based on (a) CLEC1B (b) CLEC3A (c) CLEC3B (d) CLEC12B ( e) CTSB (f) FCN1 (g) IRF7 (h) MAPKAPK2 (I) MRC1 (j) NLRP9 (k) NLRP10 (l) RIPK2 (m) SARM1 (n) TLR4 ( o) TNIP1

SUPPLEMENATRY S2
